# Information theoretics for the machine learning detection of functionally conserved and coordinated protein motions

**DOI:** 10.1101/2020.05.29.089003

**Authors:** Gregory A. Babbitt

## Abstract

Traditional information theoretic analysis of functionally conserved binding interactions described by multiple sequence alignments are unable to provide direct insights into the underlying strength, spatial distribution, and coordination of the biophysical motions that govern protein binding interactions during signaling and regulatory function. However, molecular dynamic (MD) simulations of proteins in bound vs. unbound conformational states can allow for the combined application of machine learning classification and information theory towards many problems posed by comparative protein dynamics. After both bound and unbound protein dynamic states are adequately sampled in MD software, they can be employed as a comparative training set for a binary classifier capable of discerning the complex dynamical consequences of protein binding interactions with DNA or other proteins. The statistical validation of the learner on MD simulations of homologs can be used to assess its ability to recognize functional protein motions that are conserved over evolutionary time scales. Regions of proteins with functionally conserved dynamics are identifiable by their ability to induce significant correlations in local learning performance across homologous MD simulations. Through case studies of Rbp subunit 4/7 interaction in RNA Pol II and DNA-protein interactions of TATA binding protein, we demonstrate this method of detecting functionally conserved protein dynamics. We also demonstrate how the concepts of relative entropy (i.e. information gain) and mutual information applied to the binary classification states of MD simulations can be used to compare the impacts of molecular variation on conserved dynamics and to identify coordinated motions involved in dynamic interactions across sites.

## Introduction

All variety of functions of living organisms are defined by dynamic processes that proceed over a vast multitude of size and time scales. At the systems level, a biological organism can be defined by a dynamic network of genetic and environmental interactions. Within this network are dynamic signaling pathways that form feedbacks to create temporary homeostatic stable states. In turn, each of these pathways is comprised of many dynamic molecular encounters between protein, nucleic acid and small molecule structures that bind to one another via weak and strong chemical bonding interactions. And finally, each one of these biomolecules is itself a rapidly dynamic conformation of soft matter that is thermally responsive to continuous alteration of its local solvated environment. Therefore, the study of biological function within living systems would seem to require a thorough analysis of the time series of molecular structural conformation (i.e. molecular dynamics), ultimately providing the biologist a dynamic perspective on functional molecular phenotypes. The process of organic evolution also works across these scales by first slowly accumulating chemical mutations that affect the rapid dynamics of molecular processes, and then selecting the resulting self-organized complex phenotypes that are more fit and fixing them within populations. Therefore, we should expect that when the molecular dynamics of a given protein is functionally adaptive in some way, it should produce a tendency to move in a non-random and specifically directed way, much like a machine, that is readily distinguishable from thermal noise imparted by its solvent environment. It should also preserve this characteristic motion over deep evolutionary time. Therefore, when protein sequences are functionally conserved over evolutionary time scales (i.e. millions of years), we should expect to also see molecular dynamics that are conserved over very short functional time scales (i.e. nanoseconds) as well.

Unfortunately, this necessary molecular dynamic perspective on biological function has been relegated to only a few computationally heavy fields in biology (i.e. biophysics, computational biochemistry and systems biology), and is still largely ignored by a mainstream biology that maintains a methodological focus on the comparative interpretation of static representations of biomolecular sequences, sequence annotations and protein structures housed within ever-growing online databases. While molecular biology has often been generally defined as the subset of biochemistry in which information can be substantiated and manipulated (Davies; Schrödinger and Penrose 1992; Davies et al. 2013; Walker and Davies 2013), the concept of information in biology has often been historically disconnected from its mathematical relationship with randomness in thermodynamics (Shannon 1948) and further muddied with mathematically undefined terminology like information ‘flow’, ‘transfer’ or ‘processing’ (Cobb 2013) when referring to the molecular templating involved in the replication of DNA, the transcription of DNA to RNA and the translation of RNA to protein. Fortunately one exception has been the common application of information theory to multiple alignments of binding sequences (Schneider et al. 1986; Schneider and Stephens 1990). We feel there is room to expand the mathematical formalism that already exists around this concept of information content embodied in functionally conserved biomolecular sequences to the concept of information embodied within functionally conserved motions in biomolecular dynamic simulations as well. We would go further to argue that analyses that largely ignore dynamics, rooted only upon a sequence representation of molecular biology will continue to potentially obscure a very large component of latent functional variability that connects the genotype to the phenotype through the scaling of self-organized dynamic physicochemical processes that build the complex body forms and functions that biologists define as life (Babbitt et al. 2016).

While all-atom and coarse-grained molecular dynamic studies are often used to address functional chemistry and thermodynamics of many biomolecular systems, the lack of widespread adoption of dynamic simulation by the mainstream disciplines of biology can perhaps be linked to both a lack of standard statistical analytical methods for comparing molecular dynamic simulations to each other as well as this lack of a connection between information theory and molecular dynamics as they apply to biological systems. Comparative methods are central to laboratory experimentation in molecular biology, and the information-theoretical perspective applied to DNA sequences continues to enable major advances in the fields of genomics and molecular evolution that have developed in parallel with major advances in modern sequencing technologies. Concurrent technological advances in computer hardware have also allowed larger, longer and more highly parameterized (i.e. accurate) molecular dynamic simulations (Götz et al. 2012; Salomon-Ferrer et al. 2013), and now greatly enhance our ability to investigate the link between molecular dynamics and evolutionarily-conserved function (Babbitt et al. 2014, 2016, 2018a, 2020b). However, very few methods and software currently exist allowing the exploration of function and evolution through the combination of machine learning and molecular dynamics simulation (Babbitt et al. 2018b, 2020a; Pérez et al. 2018; Wang et al. 2020). Here, we propose a generalizable method of employing machine learning-based classification of functional dynamics under the same information theoretic framework that has proven so useful for the comparative analysis of multiple-aligned protein and DNA sequences (Schneider et al. 1986; Schneider and Stephens 1990). Our method allows for the detection of functionally conserved dynamic motions using comparative molecular dynamic simulations of proteins in different functional states, without the need to resort to additional sequence-based molecular evolutionary analyses (Bakan et al. 2014). While sequence-based tests of selection can readily identify adaptively altered or functionally constrained sequences in genomes, unless they are complimented with functional studies, structural analysis, or physical simulations they generally fail to identify what exactly are the specific protein functions are the target of natural selection (Graur et al. 2013; Doolittle et al. 2014). Comparative molecular dynamic statistical methods inherently capture specific protein function that is defined by non-random machine-like motion in the protein system. When paired with information theoretics applied to molecular genetic variants, comparative molecular dynamics can potentially supply a more complete functional evolutionary picture of variation within protein space. Information theoretics applied to binary machine learning classifications of molecular dynamic simulations can also allow for identification of coordinated motions across sites caused by structural symmetries and/or allosteric interactions. Here, we first present a general computational protocol and an information theoretical framework for analyzing functionally conserved and coordinated motions of proteins by applying machine learning to molecular dynamic simulation. We then offer two case studies employing this new approach by analyzing the binding interaction of the Rbp4/7 subunits of RNA Pol II; a well-studied protein interaction involving Rbp4 N and C terminal domains, well conserved at the sequence level, and connected by a non-conserved linker region with strikingly different lengths in humans and yeast. We find a strong similarity between functionally conserved dynamic regions of Rbp4 discovered by our method, and functionally validated regions of sequence conservation reported by others (Armache et al. 2003, 2005; Sampath et al. 2003; Meka et al. 2005; Sharma et al. 2006; Zhao et al. 2012). We supplement this with an analysis of functional DNA interaction of the human TATA binding protein, highlighting a pervasive dynamic interactions that share mutual information across all sites of the protein when in its DNA bound state.

## Methods

### A practical simulation protocol for identifying local regions of proteins with functionally conserved dynamics

We outline a general protocol for combining molecular dynamics simulation software with machine learning packages for general programming languages with a goal of detecting functionally conserved protein dynamics related to functional binding interactions. Note: this is also the protocol implemented in our most recent software release (Babbitt et al. 2020a). Throughout the rest of the article we abbreviate this protocol as FCDA (Functionally Conserved Dynamic Analysis)

A. Utilizing one of several popular molecular dynamic (MD) simulation software suites (e.g. Amber, NaMD, Charm or OpenMM), generate two large ensembles of MD production runs for training one or more machine learning classifiers. Each ensemble should represent a functional state of the molecular system under investigation (e.g. bound vs unbound state of a protein that interacts with a nucleic acid, another protein, or a functional small molecule ligand such as ATP). See Figure 1A. To avoid pseudo-replication and to mitigate the potential driving effects of the initial conditions of the simulation, each MD production run should start at a different point in time.
B. Extract one or more features of the dynamics for training using vector trajectory analysis software such as ptraj or cpptraj (Roe and Cheatham 2013). Features for analysis might include both directionless features representing local vibration/flexibility (e.g. root mean square atom fluctuation) and/or directional features of vectors representing dynamic changes in protein shape as the simulations progress (e.g. XYZ vector distances to a central feature of the protein or solvent box). These data should be extracted over many time slices to compose a univariate or multivariate feature vector for the chosen machine learning classification model. See equation 4 in (Babbitt et al. 2020a).
C. At single amino acid resolution, train one or more machine learning classifiers implemented in R or Python on these features (i.e. the feature vector) and calculate their overall learning performance on new MD production runs that represent one of the two training states of the simulation. NOTE: learning performance calculations and information content are inter-related as they both rely upon the probability of classification into one group versus the other. In simple terms, learning performance of algorithms will increase when there is information about the motions of functional states to be learned (Figure 1B). At this point one can validate the effectiveness of the training by conducting a new MD validation run on the functional bound state of the protein and calculate local frequencies of correct classification at each residue across the whole protein.
D. To identify local regions with significant functionally conserved dynamics, calculate the local correlation of learning performance on two or more of the MD validations conducted in step C using a sliding window (Figure 1B). The MD validations can be conducted upon identical structures if the underlying motivation of the research question is functional. Alternatively, the MD validations can be conducted upon ortholog structures of identical size (i.e. complete homology) if the underlying motivation of research question is focused upon how evolution has maintained function over time. The statistical significance of this local correlation can be used to call significance to the mapping of functionally conserved dynamics. The logic here is simple. If dynamic states of local regions are occurring randomly due only to thermal noise, then learning performance will not locally correlate across MD validation runs. However, if the local amino acid residues are inducing a non-random motion that is detectable by the learner, then a local correlation in the learning performance profile will be induced. If multiple machine learners are used a canonical correlation analysis and associated significance via Wilk’s lambda can be used within the sliding window (Härdle and Simar 2007)(Figure 1C). Multiple test correction for false discovery rate should also be employed to adjust p-values in accordance for the number of sites on the protein (Benjamini and Hochberg 1995). Thus local regions with functionally conserved dynamics can be mapped as either sequence annotations or color mapped protein structures.

**Figure 1.**
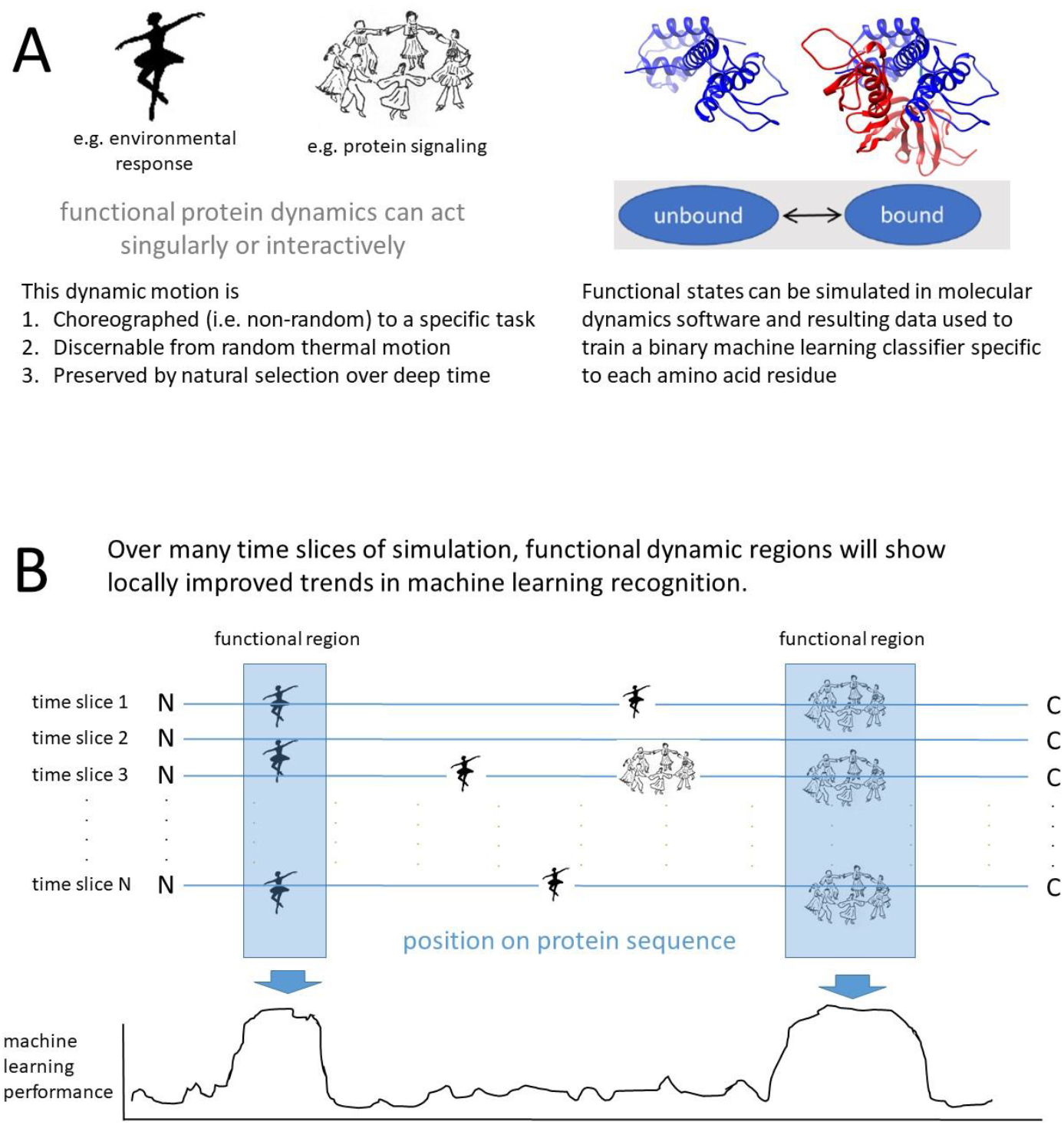
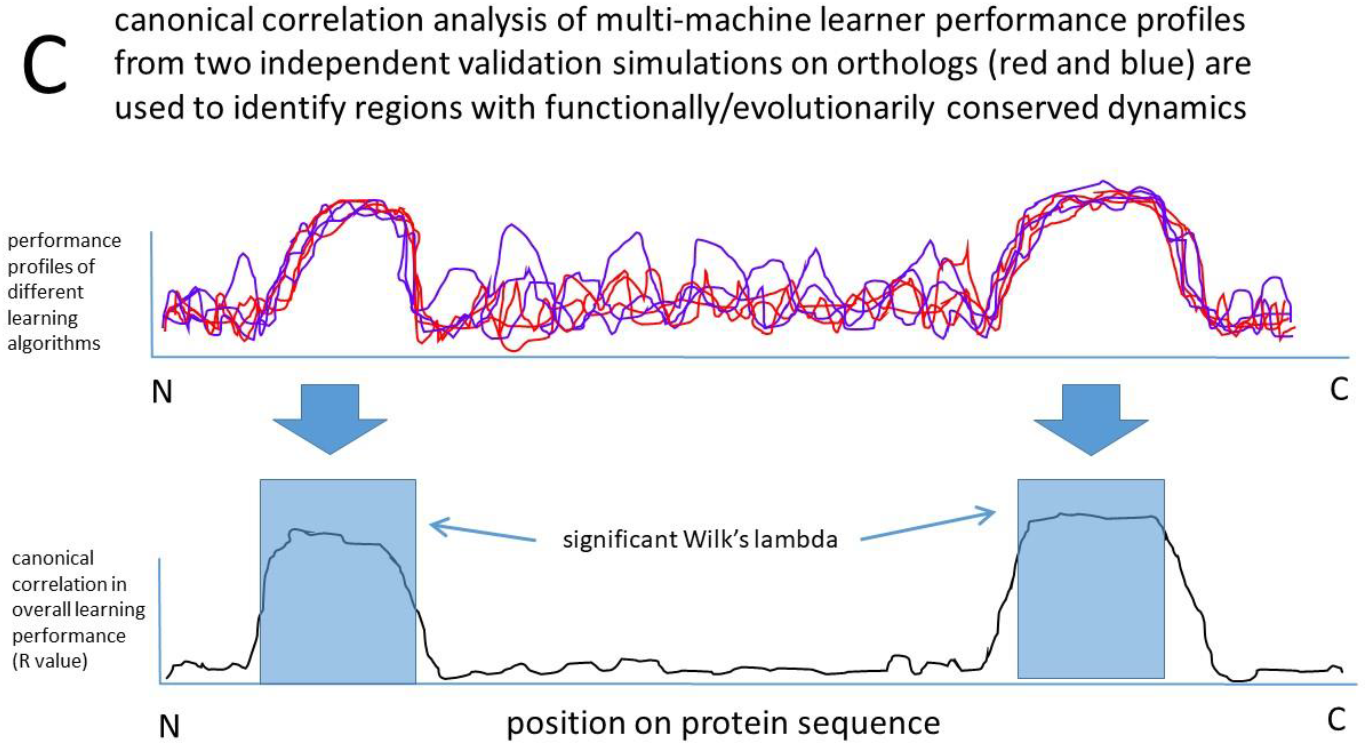
A visual analogy of Functionally Conserved Dynamic Analysis (FCDA) applied to a hypothetical ensemble of protein simulations in two functional states. (RNA POL II subunit Rbp4 bound to Rbp7 and Rbp4 in its unbound state). (A) Functional motion is evolutionarily conserved in non-random state and can be recognized by machine learning classification. (B) Local learning performance increases in functional regions of the protein and (C) can be mathematically retrieved by canonical correlation analysis of independent molecular dynamic validation runs.

### A common information theoretic framework for functional binding or protein sequences and the dynamics of their interactions

The concept of information in relation to entropy is derived from Shannon’s work ‘The mathematical theory of information’ (Shannon 1948).

Shannon Information content (I) is defined as

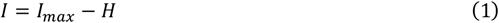

Where I_max_ is the maximum information content allowed and H is Shannon entropy observed and is defined as

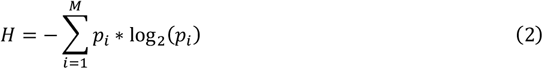

Where *p* is a set of discrete probabilities of *i* symbols from a set of possible discrete states (e.g. an alphabet) of size *M*, used to communicate a message over a discrete channel with noise (Shannon 1948).

When applied to a multiple sequence alignment of binding sites of a protein coding DNA sequence, where the frequencies of nucleobases are taken as estimates these discrete probabilities, the information content at a given site in the DNA sequence follows (Schneider et al. 1986)

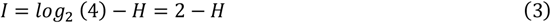

where

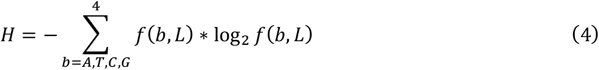

where *f*(*b*, *L*) is frequency of a given nucleobase (b) at position (L) in the best alignment or

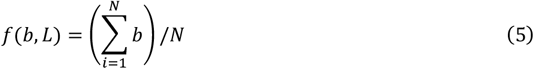

and where N is the number of aligned sequences.

This is the method that underpins the height of letter base symbols on the vertical Y axis in the well-known sequence logos plot (Schneider and Stephens 1990). And this approach can be applied to characters representing proteins by replacing the 4 possible nucleobase character states with the 20 possible amino acid character states. To obtain information for a given sequence rather than a given position within a sequence, one can average the information gained (i.e. entropy lost) by the best alignment over the whole sequence, assuming the frequencies of the nucleobases occurring at sites are independent. Therefore, the information content represents the total entropy lost as the best alignment of the sequences representing the functional site is obtained. Regions of distantly related genomes with high functional conservation will maintain high information content when aligned, as these regions are where purifying selection has not allowed random nucleobase changes to accumulate over deep evolutionary time scales.

Now if information can be imagined within the context of biomolecular sequences that represent a binding interaction, we need to now ask whether we can imagine something similar in the context of molecular motion, or dynamics, over shorter time scales as well. In many ways, the notion of a molecular form changing over time is much more amenable to Shannon’s original mathematical conceptualization of communication signal that also plays out over a relatively short time and must be distinguished from random noise. In our case, the motions of protein can be also defined in terms of ‘signal’ represented by the repeated micro-machine like motions that directly relate to a protein binding function, and ‘noise’ represented by random thermal motions caused by the continuous random collision events of solvent molecules hitting the protein. Protein function is most often defined by its binding interactions with nucleic acids, signaling molecules or other proteins, so if we could watch the molecular dynamics of a protein in two functional states (e.g. bound and unbound to a particular binding partner in a signal transduction pathway), and later apply machine learning to identify these functional dynamic states when the protein dynamics is restarted from different initial or novel conditions, then we might define the local information content in a functional dynamic context for the motion of a given residue on the protein as

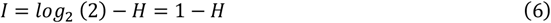

where

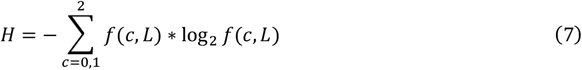

and where f(c,L) is the frequency of binary classification (c) assigned by a machine learner properly trained to recognize the dynamics of the functional state (e.g. 1 = bound, 0 = unbound) at residue location (L) taken over (T) a set of discrete time slice intervals through the total length of the molecular dynamic simulation. Therefore, the frequency of classification returned upon the deployment of a learner on a molecular dynamic simulation is

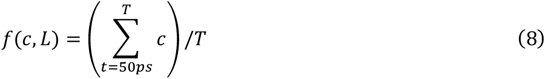

where t is the length of any given time slice interval (50 picoseconds by default) and T is the total number of time slice intervals in the simulation. An average frequency of 0.5 (i.e. the mid-point between 0 and 1), is therefore indicative if either a poorly trained learner or a lack of local functional dynamic information to be learned from a given molecular dynamic simulation.

Modern supervised machine learning algorithms are designed to derive classification states when trained on adequate amounts of real data, and molecular dynamics simulations are capable of producing large amounts of data on simulated motion due to its necessity to take very small step sizes. Hypothetically, if one applies any proper machine learning method of classification, trained on the simulations of functional dynamic states of interest, one can determine local information regarding functionally conserved protein dynamics without resorting to traditional sequence-based bioinformatics. Note that because the training classes are completely specified, the term *f*(*c*, *L*) is equivalent to a traditional measure of machine learning performance where the sum of true positive and true negative classifications divided by the sum of all total classifications (i.e. true positive/negative and false positive/negative). It is important to also note, that learners should be applied independently to the dynamics of individual amino acids as it is at this resolution of single amino acid replacement, that purifying selection will act over deep time.

### Mutual information between local learning performance profiles on independent identical simulations can identify regions of proteins with significantly functionally conserved dynamics

Assuming a machine learner is adequately trained on each amino acid to successfully discern local differences in functional dynamics states (i.e. when the protein is bound to a functional partner or not), then we can imagine running new simulation repeatedly in one of the two functional states (i.e. bound or unbound) and assessing whether local peaks in learning performance are correlated in their position along the protein backbone as opposed to just random in position. If local molecular dynamic motions of atoms are not functional as defined by the learner, there should be no local positional peaks in the learning performance or f(c,L) (Figure 1C). In this situation the average frequency of classification should be 0.5. Random thermal differences and random sampling events in new identically prepared simulations will not allow independent learning profiles collected from the same sort of simulation to be correlated. But if non-random functional motions are present on the protein backbone and the local learners can classify them according to functional states upon which the were trained, then only sampling errors will create local differences in learning performance causing correlated profiles in subsequent validation runs (Figure 1D). When multiple machine learning methods are used to reduce any artifacts caused by the sensitivity of a particular method of machine learning classification, then a multivariate or canonical correlation analysis (CCA) is required. High local R values from the CCA, generating Wilk’s lambda with a significant p-value can be used to call or map regions with significantly high local non-random dynamics associated with function (in most cases a specific binding interaction). Given two vectors representing classification states (0,1) called from *n* different machine learning methods where X = {x_1_, x_2_, …x_n_} and Y = {y_1_, y_2_, … y_n_} and new variables U and V defined via linear combinations of X and Y, the CCA seeks to find the highest canonical correlation (CC) by maximizing

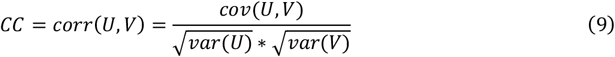

One can see great similarity of eqn 9 to mutual information in eqn. 16. Thus the CCA essentially defines the local mutual behavior of homologous amino acid residue dynamics between two MD validation runs. From a functional evolutionary perspective, this mutual dynamic behavior identified by the learner can only arise from non-random or molecular machine-like motion that is recurrent in the time scale of the simulation. It can be implied that the ability of a protein to conduct this motion must be also conserved over deep evolutionary time scales as well. In summary, the machine learning performance of a binary classification algorithm, well trained on the functionally bound and unbound dynamic states of a given protein, can be utilized to identify functionally conserved protein dynamics in both time and space. And, because the output of this analysis is binary (i.e. 0 and 1), information theory can be subsequently applied to address many interesting problems that involve comparative differences in dynamics across sites or states, as well as the coordination of dynamic states across distances define by the protein’s structure.

### Other applications of information theory to molecular dynamics

The information theoretic concept of relative entropy (also called discrimination information, information gain, or Kullback-Leibler divergence) (Kullback and Leibler 1951) can be very useful when applied to comparative questions about molecular dynamics in different functional or genetic states. In essence, the relative entropy can be conceptualized as a distance measure between two distributions of interest that is sensitive to the difference in mean, spread and shape between distributions. It can be applied to direct pairwise comparisons of the distributions of the dynamic motions themselves, or alternatively it can be applied to comparisons of local correlations in machine learning performance (i.e. information content) in validation MD runs in order to assess the impacts of molecular variation on functionally conserved dynamics. This molecular variation can represent different genetic mutations, chemical epigenetic alterations variations in small molecule binding interaction (i.e. any molecular variation of interest in the simulation of a solvated protein system).

#### A) Relative entropy between local protein motions related to functional dynamic states

The Kullback-Leibler divergence or relative entropy between two discrete probability distributions P and Q on probability space X is given by

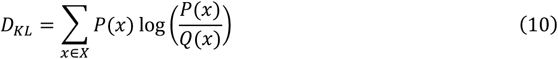

The symmetric form is the Jenson-Shannon divergence which defines the similarity between the two distributions

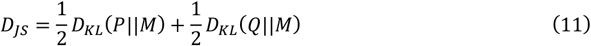

where M = ½(P+Q)

The relative entropy (RE) or similarity between the root mean square fluctuation (rmsf) of two homologous atoms moving in two molecular dynamic simulations representing a functional binding interaction (i.e. where 0 = unbound state and 1 = bound state) can similarly be described by

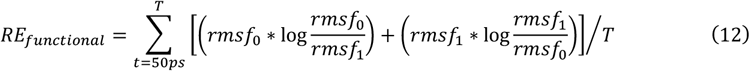

where *rmsf* represents the average root mean square deviation of a given atom over time t. More specifically, the *rmsf* is a directionless root mean square fluctuation sampled over an ensemble of MD runs with similar time slice intervals. Because mutational events in evolution most often replace entire residues, this calculation is more useful if applied to resolution of single amino acids rather than single atoms. Because only the 4 protein backbone atoms (N, Cα, C and O) are homologous between residues, the R groups or side chains are ignored in the calculation and the following equation can be applied. Because the sidechain atoms always attach to this backbone, *rmsf* still indirectly samples the dynamic effect of amino acid sidechain replacement as they are still present in the simulation.

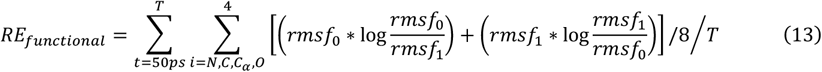

When applied to the entire protein and standardized for protein size, relative entropy can be calculated as

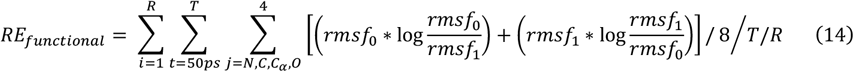

where R is the number of amino acid residues in the protein.

#### B) Relative entropy between conserved dynamic states of variants

When experimenting with *in silico* site-directed mutagenesis or with differently docked small molecule ligands, it may be more useful to compare the impacts of these variants on the functionally conserved dynamics (defined previously) rather than features like *rmsf* representing on the total dynamics. This is important as not all dynamics may be related to the protein function. The relative entropy between the local self-correlation in learning performance (i.e. functional dynamic information contributing to learning performance of the two validation runs) and each variants correlation to the validation run can prove useful for comparing the functional dynamic impacts of each molecular variant.

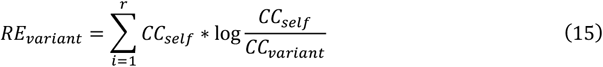

Where CC is a canonical correlation analysis of learning performance profiles along length of the protein comparing two identical MD validation runs (i.e. CC_self_) or a variant to one of the validations (i.e. CC_variant_). Empirical p-values for determining if genetic variants or drug classes differ significantly can be easily generated by randomization tests conducted with bootstrapped frequencies of classification (i.e. learning performance) generated by the time slices of MD variant runs.

#### C) Mutual information and coordinated dynamics

Mutual information (or trans-information) (Shannon 1948; Kullback and Leibler 1951) is defined as the relative entropy between the joint distribution or co-occurrence of two events A and B and the product distributions of their individual occurrence. Thus

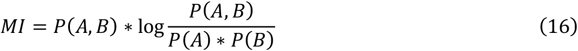

where P is the probability of two events A and B that may or may not occur independently

If we were to consider the co-occurrence of the functional dynamic classification states over time on any two sites of the protein, we can use mutual information to identify concerted dynamic functions between functional protein sites. This can potentially be very useful for the study of dynamics of proteins with bilateral binding symmetry (Matsunaga et al. 2012; Schulze et al. 2014) or allosteric regulation (Motlagh et al. 2014). If the dynamics of two neighboring sites or even two distant sites are locked together in time, they will exhibit higher degrees of mutual information. Therefore, following the machine learning-based functional dynamic analysis outlined above, a mutual information (MI) matrix calculated over all functionally conserved sites can identify regions where neighboring and/or more distant sites exert influence over each other’s dynamic function.

The mutual information between two functional dynamic classification states (0=unbound and 1=bound) at any two given residue sites (r1 and r2) calculated over t time intervals can be calculated as

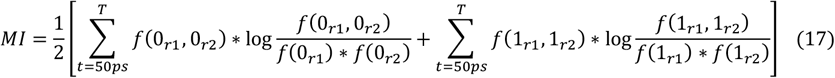

### A case study of the functionally conserved dynamic analysis (FCDA) of the RNA polymerase II Rbp4 subunit

Here we demonstrate the efficacy of our proposed method of functionally conserved dynamic analysis (FCDA), using molecular dynamic simulations of the bound and unbound states of Rbd4, the fourth largest subunit of RNA polymerase II. Rbd4 functionally binds Rdb7 within the RNA pol II structure (Figure 2A) and has well characterized conserved regions in the N and C terminus connected by a non-conserved linker region (Figure 2B). Functional studies in yeast mutants have established clear links between many functional yeast phenotypes and deletions in these regions. We used our FCDA protocol above to examine whether functionally conserved dynamics would be associated with the functionally conserved sequence regions previously established by the work of (Sampath et al. 2003). For comparison, we compared this protein-protein interaction to a DNA-protein interaction in the form of human TATA binding protein (TBP) bound to DNA (Nikolov et al. 1996).

**Figure 2.**
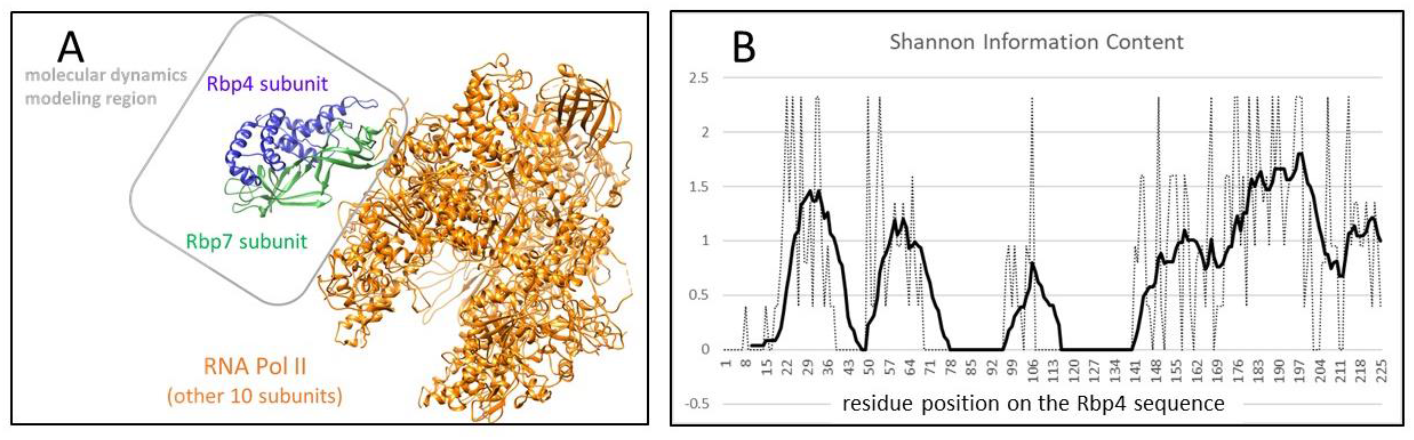
The functional binding of Rbp4 within RNA Pol II. (A) The 12 unit RNA Pol II (PDB: 1wcm) highlighting the relative positions of subunits Rbp4 and Rbp7 (PDB: 1y14) where the comparative molecular dynamics analysis of Rbp4 implemented. (B) The 10 period smoothed Shannon information content for the multiple sequence alignment of Rbp4 from Sampath et al. 2003. The alignment includes Saccharomyces cerevisiae, Schizosaccharomyces pombe, Arabidopsis thaliana, Drosophila melanogaster, and Homo sapiens.

### PDB structure preparation and MD simulation protocols

Structures of the human and yeast Rbp4 bound to Rbp7 were obtained from the Protein Data Bank (PDB). These were PDB ID: 2c35 and 1y14 respectively (Armache et al. 2003, 2005; Meka et al. 2005). We similarly obtained human TBP structure (PDB: 1cdw). Large ensembles of graphic processing unit (GPU) accelerated molecular dynamic simulations were prepared and conducted using the particle mesh Ewald method employed by pmemd.cuda in Amber18 (Case et al. 2005; Salomon-Ferrer et al. 2013) via the DROIDS v3.8 interface (<u>D</u>etecting <u>R</u>elative <u>O</u>utlier <u>I</u>mpacts in <u>D</u>ynamic <u>S</u>imulation) (Babbitt et al. 2018b, 2020a). Simulations were run on a Linux Mint 19 operating system mounting two Nvidia Titan Xp graphics processors. Explicitly solvated protein systems were prepared using tLeAP (Ambertools18) using the ff14SB protein force field (Maier et al. 2015). For TBP, we also loaded the DNA.OL15 force field (Dans et al. 2017). Solvation was generated using the Tip3P water model (Mao and Zhang 2012) in a 12nm octahedral water box and subsequent charge neutralization with Na+ and Cl-ions. After energy minimization, heating to 300K, and 10ns equilibration, an ensemble of 200 MD production runs each lasting 0.5 ns of time were created for both bound and unbound Rbp4 in both the yeast and human MD analysis. Each MD production run was preceded by a single random length short spacing run selected from a range of 0 to 0.25ns to mitigate the effect of chaos (i.e. sensitively to initial conditions) in the driving ensemble differences in the MD production runs (Babbitt et al. 2020a). All MD was conducted using an Andersen thermostat (Andersen 1980) under constant pressure of one atmosphere. Root mean square atom fluctuations (*rmsf*) were calculated using the atomicfluct function in CPPTRAJ (Roe and Cheatham 2013). All color maps on protein structures were produced in UCSF Chimera (Pettersen et al. 2004).

### Functionally Conserved Dynamic Analysis (FCDA) of bound and unbound states protein subunits within RNA Polymerase II and DNA-bound and unbound states of TATA binding protein

The signed symmetric relative entropy between the distributions of atom fluctuation (i.e. root mean square fluctuation or *rmsf* taken from 0.01 ns time slices of total MD simulation time) on bound and unbound protein states were computed using our program DROIDS and color mapped to the protein backbone with individual amino acid resolution to the bound structures using a temperature scale (i.e. where red is hot or amplified fluctuation and blue is cold or dampened fluctuation). The reference state of the protein is unbound while the query state is bound. Therefore, this pairwise comparison is used to represent the functional impact of Rbp7 binding on the Rbp4 protein’s normal unbound motion, where it is expected that contacts would typically dampen the fluctuation of atoms around the binding interface to measurable degree. In the case of TATA binding protein (TBP), the functional dynamic comparison was comprised of DNA bound to TBP and unbound TBP, again where it is expected that DNA binding should dampen the atom fluctuation to some degree. Functionally conserved dynamics were determined using the method outlined previously with a stacked machine learning model that included 7 classification methods to classify *rmsf* time series values into either ‘bound’ or ‘unbound’ classifications. The classifiers included K nearest neighbors, naïve Bayes, linear discriminant function, quadratic discriminant function, support vector machine, random forest and adaptive boosting. Significant canonical correlations in the 7 local machine learner performance profiles were determined using Wilk’s lambda and color mapped to dark gray regions on the bound Rbp4 and TBP structures. A mutual information matrix to map coordinated classifications of dynamic states over time between all amino acid residue pairs was also calculated. All software used to produce all statistical and graphical output is part of a comprehensive package DROIDS/maxDemon version 3.8 available at GitHub (https://github.com/gbabbitt/DROIDS-3.0-comparative-protein-dynamics) and our software website (https://people.rit.edu/gabsbi/) and its domain (http://proteindynamics.net). The data used to generate all the figures is deposited at https://zenodo.org/record/3820675#.XtZiRDpKhPY.

## Results

### FCDA case study in RNA Polymerase II

The relative entropy in residue backbone *rmsf* between Rbp7 bound Rbp4 and unbound Rbp4 dynamic states (i.e. eqn. 13) revealed common patterns of dampening in atom fluctuation (i.e. rapid harmonic motion) in both human and yeast. In accordance with the prediction from the functional work of Sampath et al (2003), both of our *in silico* comparative dynamics experiments show more functional dampening of atom motion in the conserved N and C terminal regions with less dampened motion in the non-conserved central linker regions (Figure 3). Note that the linker region is much larger in yeast, and our patterns of dampened atom fluctuation reflect this as well (Figure 3B). The most dampened motion coincides consistently with Rbp4’s contact surface with Rbp7 (Figure 4) which proportionally comprises a much larger fraction of the protein in human Rbp4 (Figure 4A-C) than yeast (Figure 4 D-F) and best compared in the side profiles (Figure 4B and 4E). The machine learning profiles (i.e. frequency term in eqn. 8) on the Rbp7 bound Rbp4 MD validation runs (Figure 5A/5C) indicate that motions indicative of functional binding were very well classified (i.e. recognized by the learning model) in the same regions showing more dampened motion or large negative relative entropy in Figures 3. This indicates that our machine learning classifier was able to recognized differences in *rmsf* related to functional binding between the two subunits of RNA Pol II. Significant Wilk’s lambda for canonical correlations indicate regions of functionally conserved dynamics that correspond very well with regions of dampened *rmsf* upon binding (Figure 3) and the functionally conserved regions associated with the Rbp4/7 interface identified by (Sampath et al. 2003)(Figure 5B and 5 D). The yeast linker region which is largely nonconserved at the sequence level according to Sampath (Sampath et al. 2003) shows very little conserved dynamics according to our method (Figure 5D). The human Rbp4, which lacks the large linker region, exhibits much more conserved dynamics throughout the protein (Figure 5B). The mapping of functionally significantly conserved dynamic regions to the Rbp4 structures also indicates that Rbp4 linker regions more distant from the Rbp7 interface are far less likely to be classified by learners as having a role in functional binding than Rbp4 region interacting more directly with the binding interface (Figure 6). Significance tests on the relative entropy of *rmsf* and R values derived from canonical correlations of machine learning profiles correspond roughly with the 10 period moving average of sequence-based Shannon information (r = 0.487/0.484 for human/yeast Rbp4) indicating that much of the Rbp4 protein sequence information maintained across distant taxonomic groups is contributing to the maintenance of functionally conserved dynamic motion related to Rbp7 binding (Supplemental Figure 1). Lastly, mutual information matrices (eqn. 17, Figure 7) indicate that time coordinated dynamic motions between amino acid residues are quite prevalent in the non-conserved linker regions of Rbp4, and that this is more readily apparent in the central linker region of yeast Rbp4 (Figure 7C) than in human Rbp4 (Figure 7B) or random control (Figure 7A).

**Figure 3.**
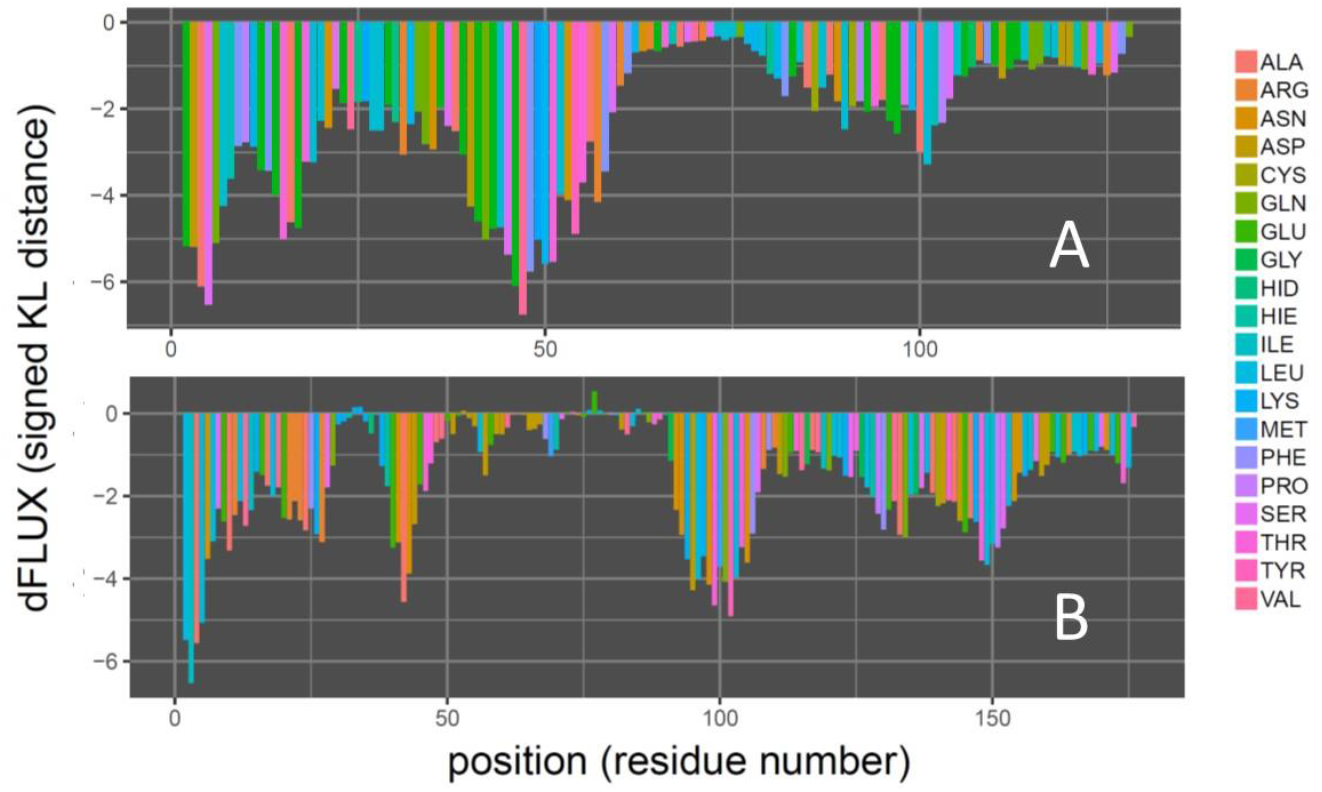
Quantifying the functional binding interaction of Rbp4/7 within RNA Pol II with a signed symmetric Kullback-Leibler distance (i.e. relative entropy of eqn. 13) comparing root mean square fluctuation (rmsf) of protein backbone for Rbp7 bound Rbp4 and unbound Rbp4 subunits of RNA Pol II in both (A) *Homo sapiens* and (B) yeast *Saccharomyces cerevisiae*. Note that more negative KL divergences or dFLUX indicates a stronger functional binding interaction between Rbp7 and Rbp4.

**Figure 4.**
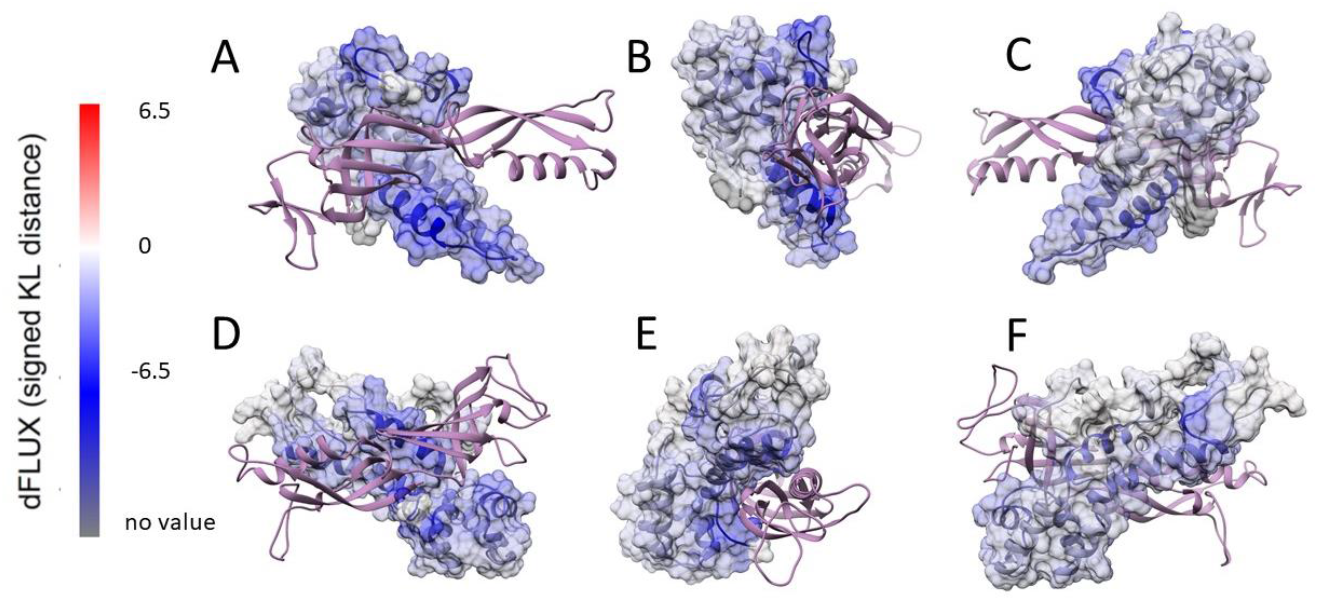
Imaging the functional binding interaction of Rbp4/7 within RNA Pol II with color mapping of signed symmetric Kullback-Leibler distance (i.e. relative entropy of eqn. 13) from Figure 3 in (A-C) front, side and back profiles of human Rbp4 and (D-F) the identical profiles for yeast, *S cerevisiae*. Note: Rbp4 is the dFLUX surface mapped structure while Rbp7 is indicated with lavender ribbon.

**Figure 5.**
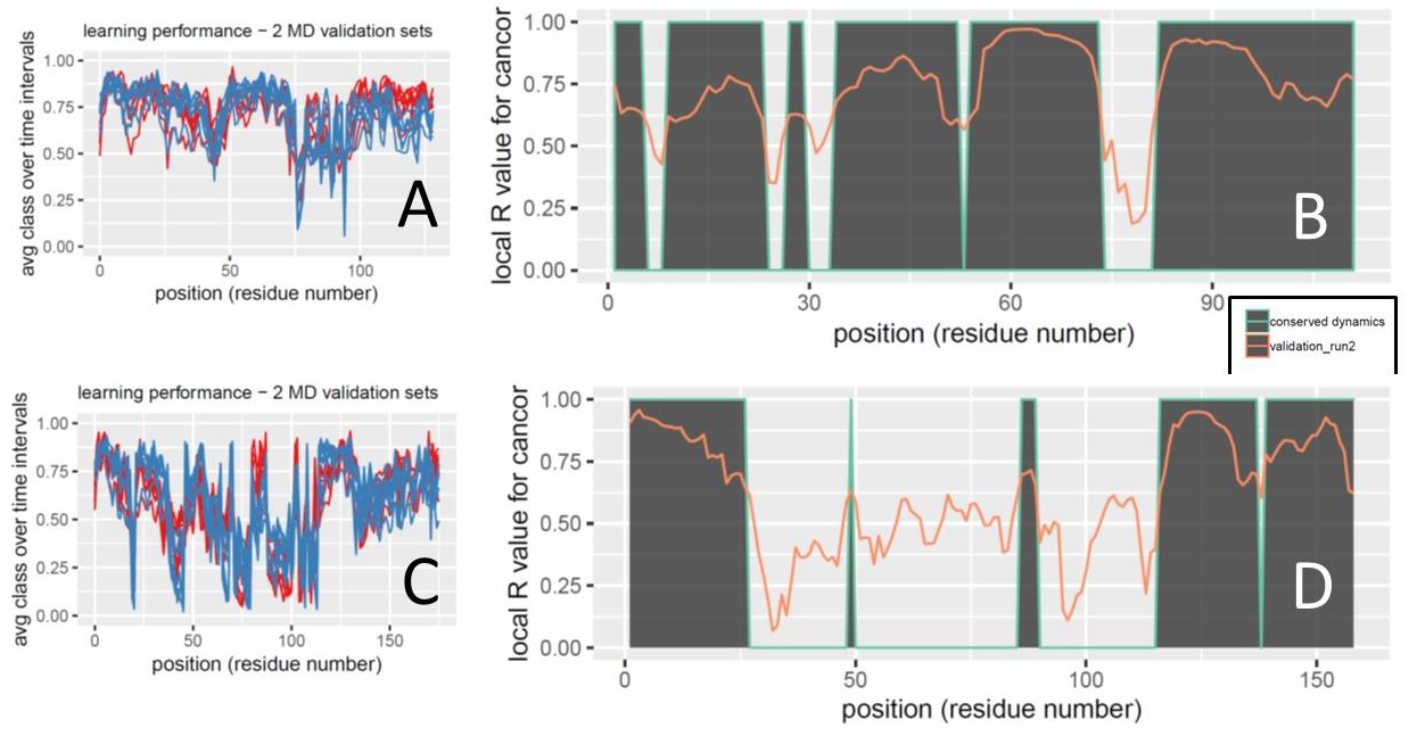
Functionally conserved dynamics of Rbp4 identified via local canonical correlation analysis of multi-machine learning performance profiles in (A-B) humans and (C-D) yeast, *S. cerevisiae*. Machine learning performance profiles (frequency term in eqn. 8) were generated in two independent molecular dynamics validation runs on Rbp7 bound Rbp4 (red and blue lines) using multiple classification methods including K-nearest neighbor, naïve Bayes, linear discriminant analysis, quadratic discriminant analysis, support vector machine, random forest and adaptive boosting (A and C). The local R value for the canonical correlation is shown over an 18 residue sliding window (orange line in B and D) and its regions with significant correlation (i.e. determined via Wilk’s lambda) is demarcated using gray shading under the green line.

**Figure 6.**
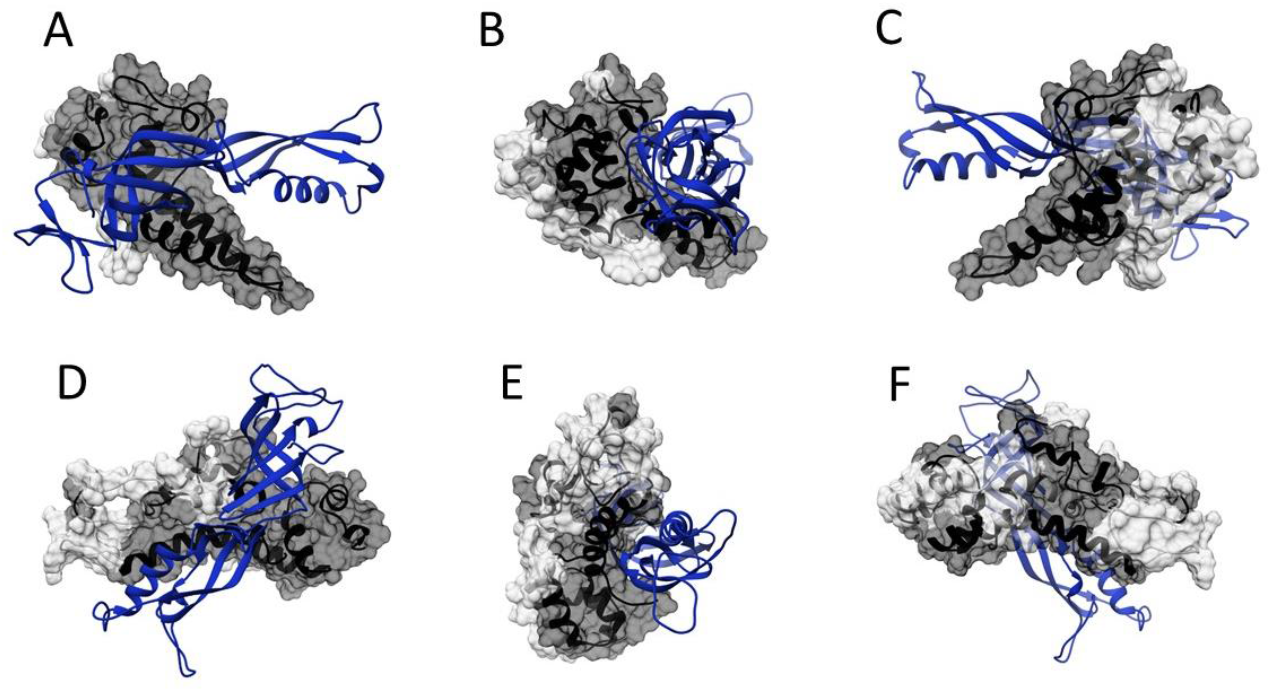
Mapping of functionally conserved dynamics of Rbp4 identified via local canonical correlation analysis of multi-machine learning performance profiles in (A-C) humans and (D-F) yeast, *S. cerevisiae*. Local regions of Rbp4 with significant functionally conserved dynamic interaction (eqn. 9) with Rbp7 (blue ribbon) are mapped in dark gray.

**Figure 7.**
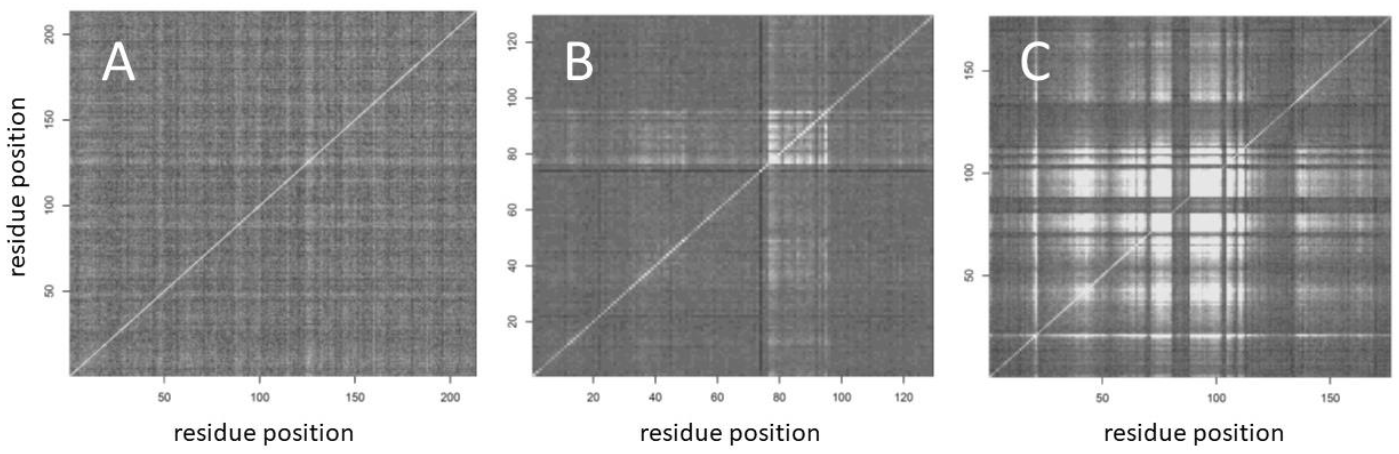
Mutual information matrices for (A) a null example with very little dynamic differences to learn and examples of Rbp7 bound Rbp4 in (B) humans and (C) yeast, *S. cerevisiae*. Black tiles indicate no matching of classification of functional dynamic states between two given amino acid residues (i.e. no mutual information from equation 17) while white indicates complete matching of dynamic states (i.e. total mutual information from equation 17). The machine learning classifier in these examples was a parameter tuned support vector machine with polynomial kernel.

### FCDA case study in TATA binding protein

We conducted an identical FCDA analysis comparing the dynamics of DNA bound TBP to its unbound form (Figure 8). Relative entropy calculations indicated where the major groove contacts of TBP are (Figure 8A and 8B). The machine learning analysis of multiple validation runs on DNA bound TBP indicated very high and significant canonical correlations in machine learning recognition of binding state across almost the entire structure of TBP (Figure 8C). A mutual information matrix also indicated very high levels of time coincidental classifications of indicating very strong coordination of functional dynamic states across the whole structure of TBP (Figure 8D). As the DNA binding partner of TBP is much more rigid than the Rbp7 protein binding partner to Rbp4, there is consequently much more mutual information between the functional classifications of dynamics between different protein sites.

**Figure 8.**
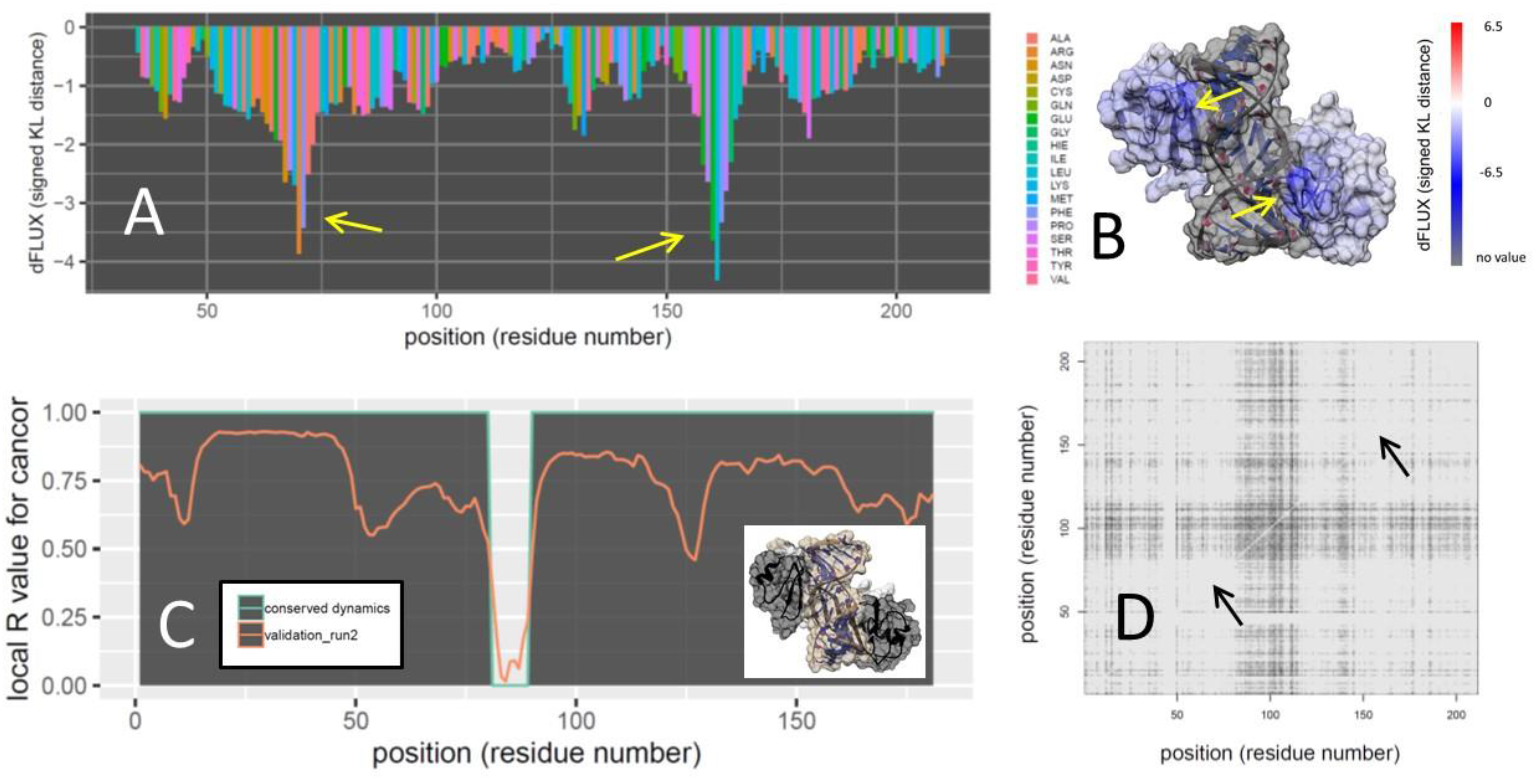
Functionally conserved dynamic analysis (FCDA) and the associated information theoretics for human TATA binding protein in DNA-bound and unbound functional states (PDB: 1cdw) showing (A) negative relative entropy associated with binding that is also (B) color mapped to the DNA bound protein structure. Note that blue regions indicate dampened protein atom fluctuations associated with functional binding. Yellow arrows indicate loop structures that interact with DNA major groove. (C) Functionally conserved dynamics regions are determined via significant Wilk’s lambda value (p<0.01) determined from canonical correlation analysis of machine learning validation profiles within 30 residue sliding window. Conserved dynamics are indicated by dark grey regions in both the plot and the structure (inset image). (D) A mutual information matrix indicating where functional dynamic classifications coincide over time is also shown. Lighter tiles indicate high degree of time dependent mutual information in functional dynamic classifications between amino acid pairs.

## Discussion

We have introduced an information theoretical formalism and a general protocol for adapting machine learning classification applied to molecular dynamics simulation towards the task of identifying functionally conserved and coordinated motions in proteins. While molecular dynamics simulations are traditionally computationally heavy, this method can now easily be accelerated on modern graphics processors and potentially allows the molecular biologist to step beyond the functional abstraction of information contained in sequences, and to directly analyze information content in the functional motion of proteins in accurate biophysical simulations. This allows for the definition of functional conservation within the known context of specific function of the proteins under investigation. Because machine learning classification binary digitizes molecular dynamic output, it allows information theory to be analytically applied in a variety of useful ways for discerning random thermal events (i.e. entropy) from non-random motion (i.e. function) in the context of protein interactions with other proteins, nucleic acids or small molecules. This protocol can now be implemented in the latest releases of our labs software website https://people.rit.edu/gabsbi/ or http://proteindynamics.net and published software notes (Babbitt et al. 2018b, 2020a). The information theoretics incorporated in the latest release of DROIDS+maxDemon version 3.8 include all the methods introduced here, including the comparison of *rmsf* between functional states via relative entropy, the identification of functionally conserved dynamics (FCDA) in new MD runs, the comparison of relative impacts of variants on conserved dynamics via relative entropy, and the identification of coordinated dynamic motions across sites on the protein via the calculation of a mutual information matrix.

We have demonstrated the efficacy of this information theory based machine learning approach to identifying conserved dynamics by using simulations of functional binding in the well-understood Rbp4/7 subunit interaction in RNA polymerase II. Our method can quantify differences in local dynamics using relative entropy calculations and can recognize and map regions of local conserved dynamics by identifying non-random local learning performance peaks indicative of functional training states that defined the relative entropy calculations. Most importantly, our results computationally confirm the main functionally conserved relationships of the N and C terminal regions investigated through prior sequence analysis and site-directed mutagenesis studies (Todone et al. 2001; Sampath et al. 2003; Sharma et al. 2006; Zhao et al. 2012). We also demonstrate that the pairwise calculation of mutual information can be used to identify time coordinated dynamics even in regions where sequences and binding dynamics are not conserved (e.g. linker regions). Site-directed mutagenesis studies on Rbp4 have indicated that some mutations in non-conserved regions can have functional effects on phenotype. Linker regions that are not conserved at the sequence level can still have important physical function imparted by structural rigidity. By isolating time-coordinated molecular dynamics between sites, our mutual information matrices capture this feature of linker regions very well.

Our analyses of TATA binding protein (TBP) also showed the potential effectiveness of our computational approach in illuminating important functional dynamics. First of all, the relative entropy calculation on atom fluctuation (*rmsf*) highlights the locations of major groove contacts very well, and quantifies the binding effect as much larger than the rest of the protein. Second, this analysis also indicates that the whole structure of TBP is dampened in its motion during binding even outside of these major groove contacts, and this dampening is largely a function of any given protein atom’s proximity to the DNA. The realization that nearly all of the TBP protein is functionally involved in binding is also confirmed by the machine learning performance analysis and identifies almost the whole TBP structure as exhibiting significantly functionally conserved protein dynamics. Lastly, the mutual information matrix of all pairwise amino acid residues indicates very high levels of time coincident machine classification of functional dynamics states in the validation runs, even across very distant parts of the protein. This suggests that, unlike Rbp4, which binds the softer matter of Rbp7 and in two distinct regions separated by a linker section, the much higher rigidity of DNA is likely involved in coordinating long range dynamic correlations in concerted motion across the whole TBP structure when it is in its DNA bound state. Thus, the lack of a designated non-conserved linker region in TBP, combined with the rigidity of its DNA binding partner, creates very large degree of mutual dynamics when compared to Rbp4, which only seems to show mutual dynamics in its linker region. Our method captures this aspect of shared dynamics very well.

Molecular biologists typically investigate molecular function by discovering DNA/protein sequences and then conducting a series of expensive laboratory studies that probe function via experimental manipulation of these sequence changes. While protein structures and molecular phylogenetic analyses are often characterized as part of this pipeline, very rarely is computer simulation of dynamic motion of protein structure ever employed. However, this is beginning to change as more molecular evolutionary studies are beginning to include molecular dynamics simulations in their repertoire (Saldaño et al. 2016; Biagini et al. 2018; Campbell et al. 2018; Dong et al. 2018; Johansson and Lindorff-Larsen 2018; Otten et al. 2018; Adams et al. 2020). Methods of biophysical simulation of protein dynamics have been under continuous development for many decades. However, most of the questions addressed through simulation are physical rather than comparative in nature. The lack of a common comparative method for molecular dynamics data has been problematic. The complexity and size of the data sets that these simulations produce is also quite challenging. Many reviews of best practices in molecular dynamics often warn users of the danger of comparisons based upon inadequate sampling, but seldom offer proper statistical framework to mitigate this problem (Grossfield et al. 2018; Braun et al. 2019). Recently, our lab has developed a platform for statistical comparison and machine learning applications to molecular dynamics (Babbitt et al. 2018b, 2020a). Here, we have broadened this application under an information theoretical framework to comparative dynamics that has the potential to illuminate the functional consequences of molecular changes at an all-atom resolution that cannot be achieved with traditional molecular biology. Our computational protocol helps to solve an important problem of scale in molecular biology; the investigation of functional molecular motions at the very rapid time scales and very small size scales at which it occurs. It also offers the ability to connect these very rapid short time scales to the very long time scales of molecular evolution. Frameworks where machine learning is applied to molecular dynamic simulation can allow proper investigation of this complex and largely invisible process involving time scales that are not easy for humans to directly observe.

## Acknowledgements

I thank and acknowledge Dr. Ernest P. Fokoue, Dr. Miranda Lynch and Dr. George M Thurston for very helpful conversations regarding this subject.

---

**Supplemental Figure A.**
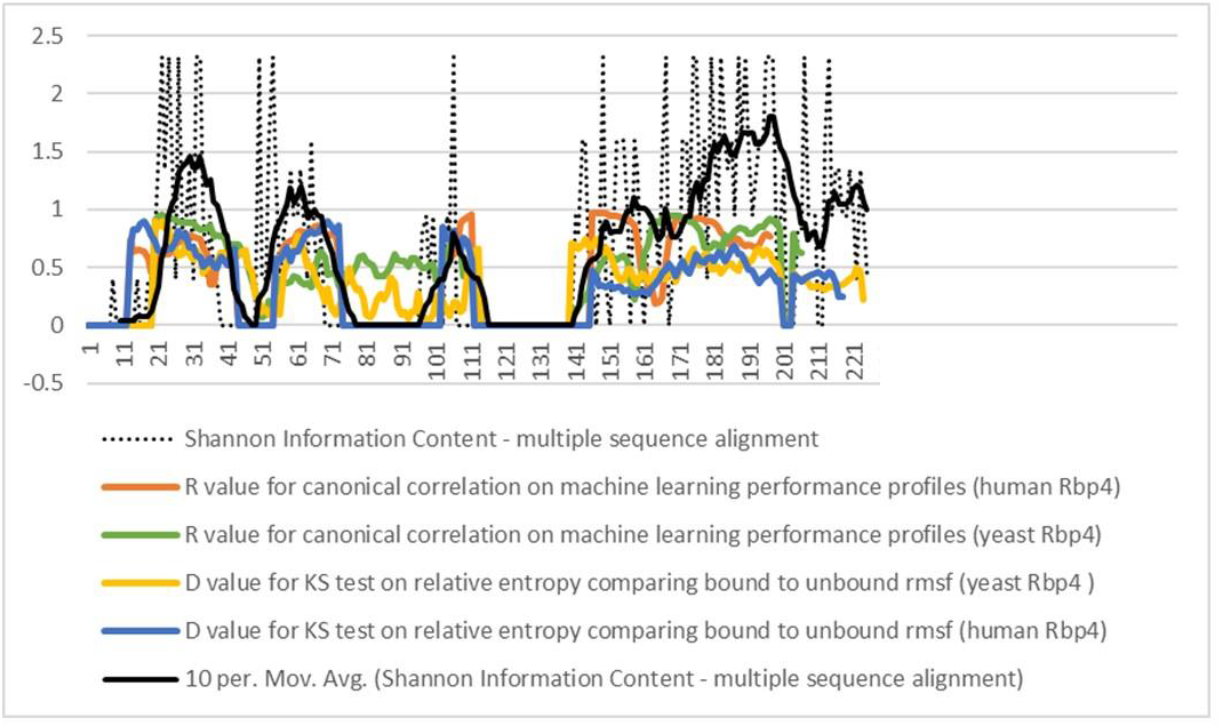
Co-occurrence of sequence-based Shannon information content and functionally conserved molecular dynamics of Rbp4/7 interaction within RNA Pol II. The Shannon information (black lines) was calculated using the multiple sequence alignment of Sampath et al. 2003. The metrics of functionally conserved dynamics (R values) and maximum statistical differences in the empirical cumulative distributions of root mean square fluctuation of Rbp4 in bound versus unbound states (D values) are shown as colored lines for comparison. Note that zero values indicate large gaps in the main alignment where yeast linker sequence is not represented in human sequence.

