## Supplementary figures and images for "Information theoretics for the machine learning detection of functionally conserved and coordinated protein motions"

### Supplemental Figure A

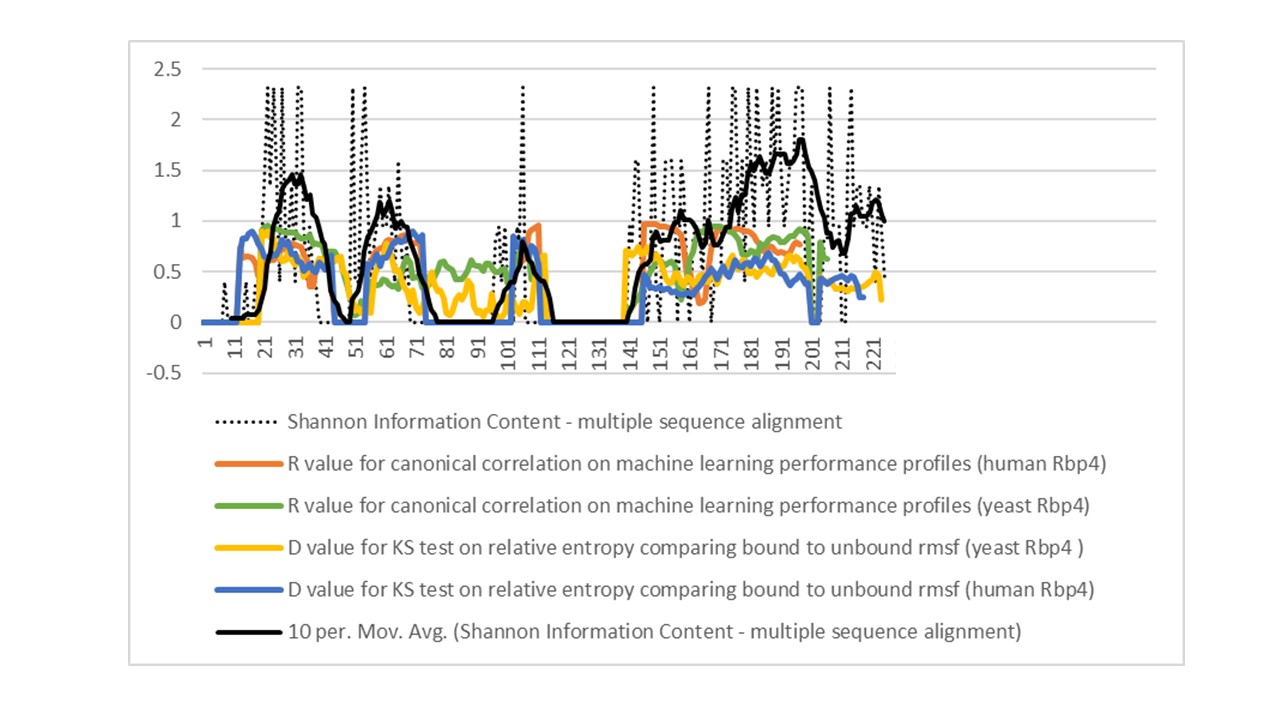
